# A simple Dot Blot Assay for population scale screening of DNA methylation

**DOI:** 10.1101/454439

**Authors:** Nelia Luviano, Sayuri Diaz-Palma, Céline Cosseau, Christoph Grunau

## Abstract

The study of epigenetic changes in natural and experimental populations has increased the need to find a cost-effective and high throughput method to analyze multiple samples to effectuate a population-wide screening to study epigenetic changes triggered by biotic or abiotic stress. One of the most studied epigenetic marks is global DNA methylation, its measurement is used as a first step to differentiate methylation between individuals. There is a wide range of methods designed to detect genome-wide 5 methyl-cytosine (5mC) that differ in sensitivity, price, level of expertise required, but as a general rule, require large amounts of DNA and are relatively expensive. This is a limit for the analysis of 5mC in a large number of individuals as a prerequisite to population-wide testing of methylation markers. In this work, we evaluated a method based on antibody recognition of 5mC to measure the DNA methylation level of individuals of the species *Biomphalaria glabrata*, the intermediate host of schistosomiases, a neglected tropical disease. We validated the method to complete a large screening in the genome of *B. glabrata* snails treated with a chemical inhibitor of DNA methylation; however, the method can be applied to any species containing 5mC. The dot blot assay is a suitable method to perform a large-scale screening of global DNA methylation to compare 5mC levels between individuals from different natural or experimental populations. The dot blot method compares favorably with methods with an equivalent sensitivity such as the Enzyme Linked Immunosorbent Assay (ELISA) kit since it requires a smaller amount of DNA (30 ng) is less expensive and allows many more samples to be analyzed.

## Introduction

Epigenetic mechanisms refer to heritable and reversible alterations in gene expression or cellular phenotype originated by changes other than modifications in the underlying DNA sequence (Nicoglou and Merlin, 2017). There are at least four carriers of epigenetic information: histone modification, non-coding RNA, location of genes in the nucleus and DNA methylation. The latter consists of the addition of a methyl group to a nucleotide, usually in the carbon 5 of the cytosine pyrimidine ring forming 5-methylcytosine (5mC). It is present in protists, plants, fungi and animals. DNA methylation is catalyzed by a family of conserved DNA methyltransferases (DNMTs). In most animals studied, DNA methylation occurs principally at CpG dinucleotides (Colot and Rossignol, 1999, Sarda et al., 2012) but methylation can occur in CHH and CHG (H= A, T, C) contexts in plants.

The types and levels of genomic DNA methylation varies significantly between species from undetectable DNA methylation (e. g. nematode *Caenorhabditis elegans*, the yeast *Schizosaccharomyces pombe* and *Saccharomyces cerevisae*) to very high levels in vertebrates (60-90% of all CpGs methylated) and most plants (Hendrich and Tweedie, 2003). The earliest methods to measure DNA methylation were based on the separation of methylated and unmethylated deoxynucleosides. One of the first techniques to measure 5-mC quantitatively was the reversed-phase high performance liquid chromatography (RP-HPLC). The quantitative measurement of DNA methylation with this method is based on the relative intensity between cytosine and 5 methylcytosine (5mC) fractions of hydrolyzed DNA (Kuo et al., 1980). HPLC was useful to compare global DNA methylation amongst different species, but has limitations (Harrison and Parle-McDermott, 2011), in particular, due to the high amount of DNA (~2.5 μg) necessary to quantify 5mC. Liquid chromatography combined with tandem mass spectrometry (LC-MS/MS) improved sensitivity and requires much smaller amounts of hydrolyzed DNA sample (50-100 ng of DNA sample) in addition, this technique is not affected by poor-quality DNA (Song et al., 2005). High performance capillary electrophoresis (HPCE) is another alternative method that is faster, low-cost and more sensitive than HPLC (Fraga et al., 2000). Nevertheless, a certain level of expertise and sophisticated equipment is necessary to perform such analyses not always available in research laboratories.

Bisulfite genomic sequencing is recognized as the gold standard method that allows for single-base resolution measurement of DNA methylation. Bisulfite treatment (Frommer et al., 1992) transforms the non-methylated cytosine into deoxy-uracil that will be read as thymine when sequenced, while 5-methylcytosine (5mC) remains intact and is still read as cytosine. Furthermore, the bisulfite genomic sequencing method works as a fundamental principle to several derived methods to quantify DNA methylation, i. e. Methylation Specific PCR (MSP), Combined Bisulfite Restriction Analysis (COBRA), and many other techniques depending on the application (Li and Tollefsbol, 2011). Complete bisulfite conversion is essential in order to have reliable quantitative methylation analysis, if the total conversion is not accomplished, unmethylatedcytosines can be mistaken for methylated residues and result in partial methylation profiles. The method requires also PCR and often sequencing and is therefore time-consuming and relatively expensive

Global DNA methylation can be quantified also by 5-methylcytosine-specific antibody combined with fluorescence staining. The analyses of quantitative DNA methylation can be accomplished by the analysis of an image with a charge-coupled device camera and sampled results need to be compared with results acquired from cells with known methylation levels (Veilleux et al., 1995). Another method to screen the global 5mC level is the Enzyme Linked Immunosorbent Assay (ELISA), there are several commercially available kits, the procedure involves the DNA manipulation in a well plate followed by consecutive incubation periods. Firstly, with a primary antibody against 5mC, then a labelled secondary antibody and finally with colorimetric/fluorometric detection reagents. However, only large variations in DNA methylation (~1.5-2 times) can be determined using this method due to the high level of inter and intra-assay variability and therefore this method is only suitable for the rough estimation of DNA methylation (Kurdyukov and Bullock, 2016).

More recently, all techniques have been applied to more technically advanced systems such as DNA cleavage by methylation-sensitive restriction enzymes combined with polymerase extension assay by Pyrosequencing, called LUminometric Methylation Assay (LUMA) (Karimi et al., 2006); or combined with genomic microarrays (Schumacher et al., 2006, Weber et al., 2005). Another system is Methylated DNA immunoprecipitation (MeDIP) that separates methylated DNA fragments by immunoprecipitation with 5mC-specific antibodies, the enriched methylated DNA can be evaluated in a genome-wide approach by comparative genomic hybridization against a sample without MeDIP enrichment (Vucic et al., 2009)

With the emergence of population-wide epigenetic screens, global DNA methylation measurements are often used as a first step to differentiate methylation between individuals. Consequently, this increases the need to find a cost-effective and high throughput method that allows analyzing multiple samples to study DNA methylation changes triggered by e.g. biotic or abiotic stress. There exists a wide range of methods designed to detect genome-wide 5mC that differ in sensitivity, price, level of expertise required, but as a rule, require either large amounts of DNA or are relatively expensive (Sant et al., 2012, Laird, 2010, Kurdyukov and Bullock, 2016). This is a limit to the analysis of 5mC in a large number of individuals as a prerequisite to population wide testing of methylation markers.

In this work, we evaluated a method based on antibody recognition of 5mC to measure the DNA methylation level of individuals of the species *Biomphalaria glabrata*. We believe that the method can be used with any methylated DNA.

*B. glabrata* is a mollusk, intermediate host of the human parasite *Schistosoma mansoni*, the causative agent of schistosomiasis, the second most severe parasitic disease in terms of morbidity just after malaria (Walker, 2011). About 2% of cytosines are methylated in the genome of *B. glabrata* (Fneich et al., 2013) and DNA methylation machinery plays probably a role in parasite-host interaction (Geyer et al., 2017). But as in most invertebrates, the precise function of 5mC in its genome remains enigmatic. We developed our method to measure changes of DNA methylation upon exposure of snail populations to chemical stress and to see if these modifications in DNA methylation produced changes in phenotypic traits (Luviano et al. manuscript in preparation).

Our method consists of immunological detection of 5mC and allows for fast screening of changes in DNA methylation in a large number of samples. It is also a simpler and less expensive screening strategy that compares favorably with methods with an equivalent level of sensitivity.

## Results

5mC methylation of DNA samples of *B. glabrata* extracted by different methods was measured. The NucleoSpin Kit improved with zirconium/silica beads method presented the most reproducible results and it was the only method that allows discriminating clearly the denaturated, the naturated and the renaturated samples (Fig.1). Since the methyl group (CH3) is located inside the double DNA helix, it can only be detected by the antibody against 5mC if DNA is properly denaturated.

**Fig.1.**
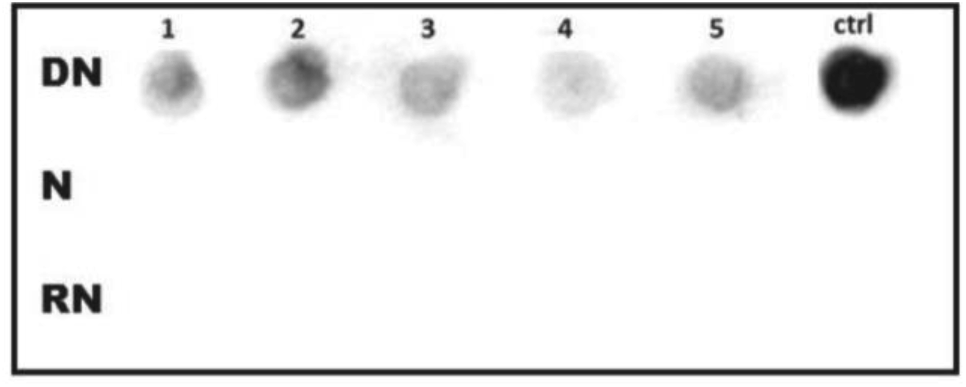
Methylated Cytosine Dot Blot of five tissue samples of *B. glabrata* obtained with the NucleoSpin kit and zirconium/silica beads method, with 120 ng of DNA. Exposure time: 35 sec. Control: HeLa DNA (200 ng). Top DN: denatured with NaOH at 42°C, middle N: non-denatured, bottom RN: renatured after 1 h at room temperature means it returns to the non-denatured form. As expected, there is no signal for the non-denatured and renatured samples.

To standardize our method, we used HeLa DNA as positive control and PCR products as negative one. Methylation level in HeLa cells is 2.3%± 0.22 of 5mC of total cytosines (Diala and Hoffman, 1982) and 0% in the case of DNA amplified in vitro by PCR and thus unmethylated. In order to test the linear range of the method, a correlation was calculated between the mean spot densitometry obtained from membrane imager and the amount of input sample 5mC in pg. The results showed a strong linearity between 5 mC amount and mean spot densitometry, therefore we decided to measure by this method a large number of *B. glabrata* samples (Fig.2). Each ng of HeLa cells contains 5.1 pg of 5mC, this value was obtained by calculating the molecular weight of 2.3% of cytosines in the human genome composed by 3.2 × 10^9^ nucleotides and a 40% GC content (Li, 2011).

**Fig.2.**
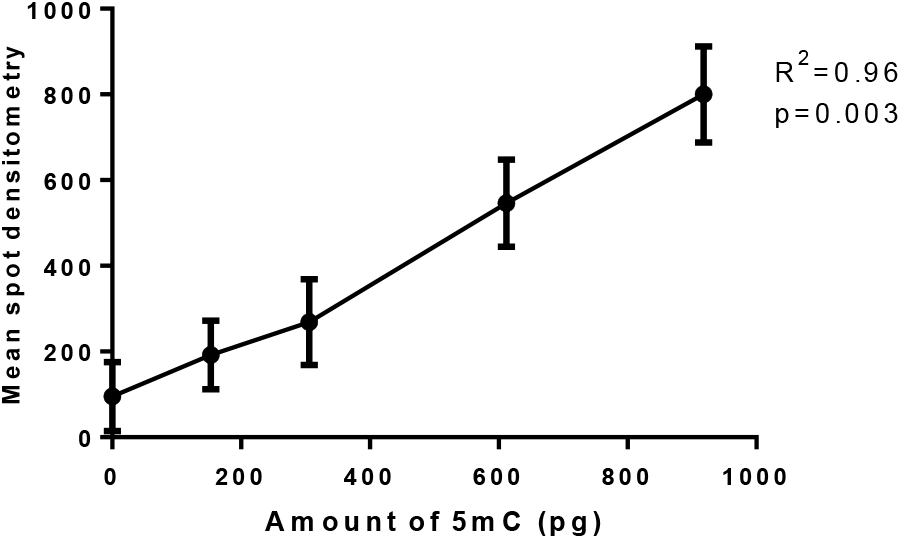
Linearity of the mean spot densitometry of the HeLa cells obtained from dot blot assay and the amount of 5mC in picograms in input sample. Each ng of HeLa cells contains 5.1 pg of 5mC, five points showed in the graphic correspond to 0, 30, 60, 120 and 180 ng of HeLa DNA

When we used as reference the densitometry/nanogram value of HeLa cells (Positive Control densitometry/nanogram value =6.92±1.2 and Positive Control 5mC%= 2.3%) to calculate by the next equation the 5mC% from *B. glabrata* samples:

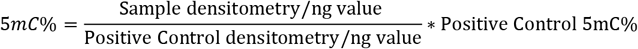

We obtained 1.97 % (Fig. 3b), which is concordant with the known methylation level in *B. glabrata* whose genome has 2% of cytosines methylated (Fneich et al. 2013). As expected, when we calculated the 5mC% in zebularine treated snails we detected a decrease in DNA methylation, the mean 5mC% decreased from 1.97% to 1.74% however, this decrease is statistically not significant (t=1.15, df=45.29, p=0.25).

**Fig.3.**
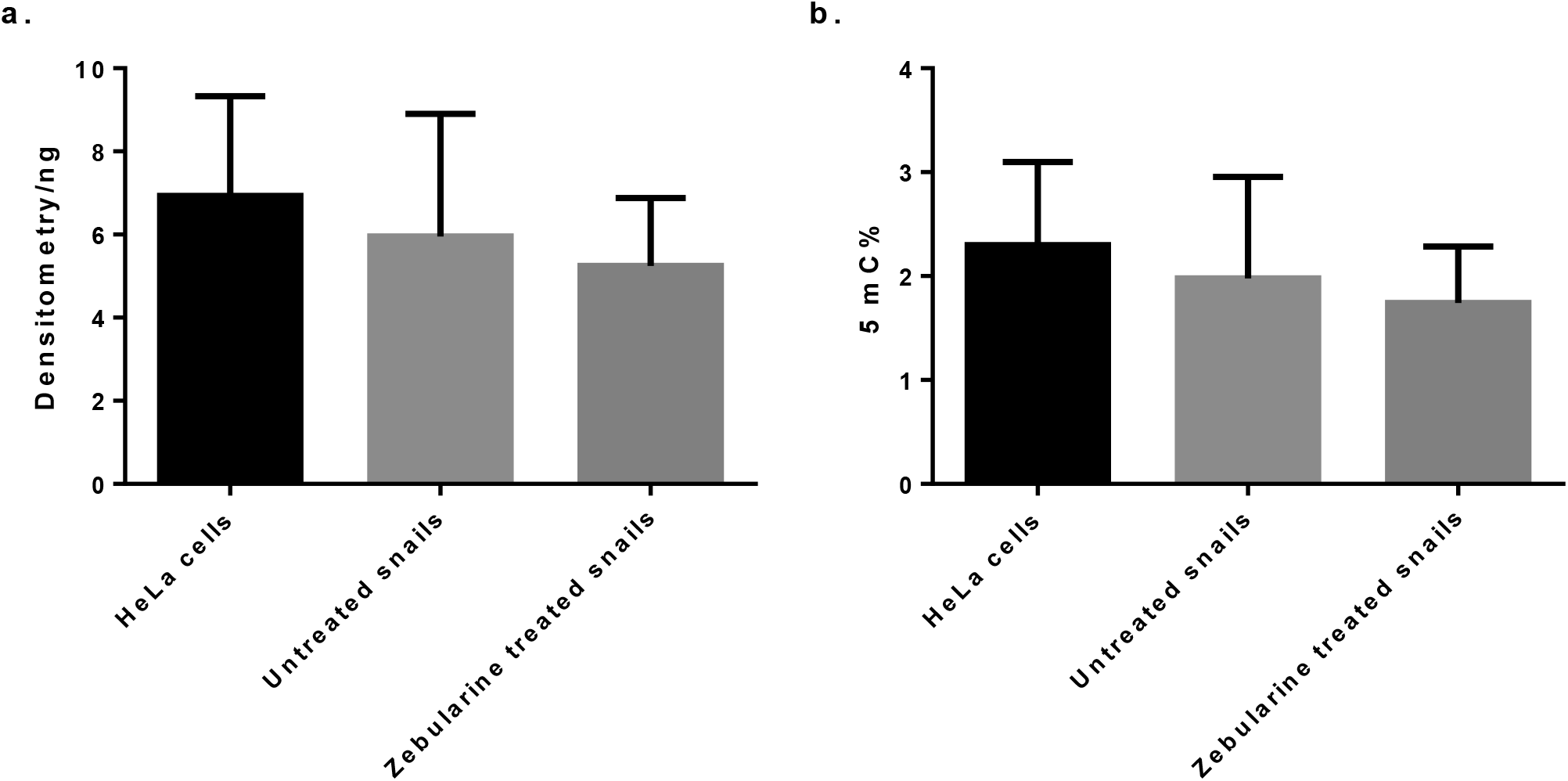
a) Densitometry/nanogram obtained by the dot blot assay of HeLa cells (x̄ =6.92±1.2), untreated snails (x̄ =5.95±0.53) and zebularine treated snails (x̄ =5.24±0.29). b) 5mC% calculated for HeLa cells (x̄ =2.3%), Untreated snails (x̄ =1.97%) and treated snails (x̄ =1.74%).

## Discussion

Three different methods to perform DNA purification before DNA methylation detection were tested in order to optimize the DNA extraction method adapted to the methylated cytosine dot blot assay. The method phenol/chloroform is widely used and requires reagents that most laboratories possess and is not expensive, however, it is a time-consuming method as it requires a lysis phase overnight. The E.Z.N.A kit is a simple, rapid and cost-effective method for the isolation of DNA, with the spin-column based technology, multiple samples can be processed in parallel. However, the overnight lysis is still required when applying this method. The NucleoSpin kit uses the same spin-column based approach as E.Z.N.A kit, but in order to improve this method zirconia/silica beads can be used to effectuate a mechanical cell lysis to speed up lysis phase comparing with other methods. In this work, the NucleoSpin kit improved with zirconia/silica beads was the method that gave us the more reproducible results. Furthermore, it was the only method that allowed us to differentiate between the denatured and non-denatured samples after dot blot assay was applied. The other two methods showed a binding of the antibody anti-5 methyl cytosine even in the non-denatured samples which is not possible since the CH3 groups of the cytosine are inside the double helix of DNA and therefore it can be only detected if we open the double-strand DNA (denaturation). For this reason, the signals from non-denatured samples were interpreted as non-specific binding of the antibody.

The exposure time suitable for our membranes in the CCD imager was 200 secs to avoid signal saturation, the exposure time can vary depending on experimental conditions but have to be always below the point of saturation of the most concentrated spot, detected by imager software. Otherwise a plot between signal and exposure time must be done in order to identify the linear range and select an exposure inside this range. Once the experimental conditions were established for the dot blot assay, we did not change the input DNA sample, membrane size, denaturation time, transfer method, transfer time, antibody solution, antibody incubation time, temperature or exposure time in all experiments as these factors can alter significantly the detection signals. All the steps in the protocol were homogeneous to avoid high intra assay-variability. In addition, positive and negative controls have to be present in all membranes to be able to compare results between them.

Dot blot method compares favorably to methods with an equivalent sensitivity like ELISA. The sensitivity of ELISA-based global DNA methylation assays vary from 0.1 ng to 10.5 ng of 5-mC DNA, our Dot Blot method showed a sensitivity of 0.15 ng, very similar to the better detection limit of ELISA. The Methylated Cytosine Dot blot assay allows for high throughput samples; it provides a measure of global 5- methylcytosine in the genome in a short period of time. In our hands, 288 samples with replica per day as multiple membranes can be incubated at the same time, and at a price (3€ per sample approximately) that compares favorably to other methods like the commercially available ELISA kit that allow to do 96 samples with replica per day if two plates are done at the same time and at 14€ per sample approximately.

After applying the methylated cytosine dot blot assay to our samples, we can conclude that it is a method adequate to perform a large-scale screening of global DNA methylation to compare 5mC levels between individuals of different natural or experimental populations. We validated the method to complete a large screening in the genome of *B. glabrata* snails treated with a chemical inhibitor of DNA methylation. This method requires a very small amount of DNA material (30-180 ng).

## Materials and Methods

### Ethics statements

*B. glabrata* Brazilian strain (Bg BRE) was used in this study. The mollusks are maintained at the IHPE laboratory facilities; they are kept in aquariums and fed with lettuce *ad libitum*. The Direction Départementale de la Cohésion Sociale et de la Protection des Populations (DDSCPP) provided the permit N°C66-136-01 to IHPE for experiments on animals. Housing, breeding and animal care were done following the national ethical requirements.

### Zebularine treatment

One hundred *B. glabrata* BRE snails (5-7 mm in size) were maintained in 1L of freshwater in the presence or absence of the demethylating agent zebularine (Sigma, France, Cat. No. 3690-10-6) at a concentration of 10 μM. The water and the fresh zebularine were replaced once at the same concentration, the replacement was performed after 3 days and 22 hours. After 10 days of exposure, the drug was removed and replaced by only water. Snails were then raised in the plastic tank during 70 days. At day 70, snails were collected, individually wrapped in aluminum sheets, and stored at −20°C.

### Optimization of DNA extraction

Three DNA extraction methods were optimized in terms of cost, scale, effectiveness and time. For this purpose, 5 samples of *B. glabrata* were isolated by phenol-chloroform method, 5 samples by the E.Z.N.A ^®^Tissue DNA Kit (Omega Bio-Tek, Ref D3396) and 5 with the NucleoSpin^®^ Tissue Kit (Macherey-Nagel, Ref #740952) combined with zirconium/silica beads.

For the phenol/chloroform method, tissue samples of *B. glabrata* were incubated overnight at 55°C with 1 ml of lysis buffer (20 mM TRIS pH 8; 1 mM EDTA; 100 mMNaCl; 0.5% SDS) and 20 μl (0.3 mg) of Proteinase K (Invitrogen, Cat. No. 11578916) The DNA of samples were extracted adding twice equal volumes of phenol/chloroform followed by 2× extractions with identical volumes of chloroform. DNA was precipitated with the same volume of isopropanol/sodium acetate. After centrifugation and washing with 1 ml of 70% ethanol, the pellet was dissolved in 200 μl of 1 mMTris/HCl, pH 8, and stored at −20°C.

Other tissue samples of *B. glabrata* were isolated using an E.Z.N.A Tissue DNA Kit (Omega Bio-Tek, Denmark), lysis, binding, washing and elution of the DNA was done according to the manufacturer’s protocol. This kit is based on proteinase K digestion overnight and in spin column-based technology.

The NucleoSpin^®^ Tissue Kit (Macherey-Nagel, Düren, Germany) combined with the use of zirconia/silica beads, a method developed to extract DNA from the pacific oyster *Crassostrea gigas* (de Lorgeril et al., 2018) was applied in tissue samples of *B. glabrata*. The zirconia/silica beads perform a mechanical cell lysis while lysis buffer acts chemically to break the cells. Briefly, for the lysis phase, samples were transferred into180 μl of lysis buffer and 25 μl (0.32 mg) of Proteinase K (Macherey-Nagel, Ref 740506.30) in 2 ml screw cap microtubes containing 100 μg of zirconia/silica beads (BioSpec, USA, Cat. No. 11079110z). This mix was vortexed. Tubes containing samples were submerged in liquid nitrogen and then shacked in a Mixer Mill (Retsch MM400) at a frequency of 30 Hz for 12 minutes. After that, incubation in a water bath at 56°C during 1 h 30 minutes was done.

After the lysis phase, the NucleoSpin^®^ Tissue Kit protocol was applied according to the manufacturer instructions for binding, washing and elution of the DNA. Elution was performed into a final volume of 100 μl in elution buffer. The samples were stored at −20°C.

DNA concentrations of all samples were quantified using a Qubit^®^ 2.0 fluorometer (Invitrogen) and a fluorescence-based Qubit™ dsDNA BR Assay Kit (Invitrogen, Q32853).

### Dot Blot assay

Once the DNA extraction method was optimized, 30 samples from the control group and 30 from the group treated with zebularine were isolated for the 5mC dot blot assay. DNA extracted from HeLa cells served as positive control and DNA of *B. glabrata* amplified by PCR was used as negative one. HeLa cells were a kind gift of Albertina De Sario INSERM U827 (IURC). A linear regression was done using the mean spot densitometry of HeLa cells obtained from the imaging system and the input of 5mC in picograms. After confirmation of our positive and negative controls, the dot blot assay was applied to the all samples of genomic DNA extracted from *B. glabrata* individuals.

#### Denaturation of DNA samples

Since the 5mC moiety that is detected by the antibody resides inside the DNA double helix and is therefore inaccessible, DNA must be denatured to expose the methylated site. The amount of DNA that allowed for optimal 5mC detection range was 30ng-180 ng in our case, and to effectuate dot blot we took 180 ng from each DNA sample. DNA was adjusted with MilliQ water to 10.8 μL in 0.2 mL tubes and 1.2 μl NaOH 3M was added (total volume= 12 μl). Tubes were incubated at 42°C for 12 minutes. After incubation, the samples were rapidly transferred by spotting each sample into the membrane of nitrocellulose (Hybond^®^). This step is important because samples can renaturate within a couple of hours. Each spot consists of 6 μl of denatured DNA so that each sample can be spotted in duplicates. The samples and their replicates must be spotted in a random way, for this purpose a paper grid template must be created with the label of each sample in the appropriate grid space. This grid template can be used to spot the samples into the nitrocellulose membrane (13 × 9 cm for 96 samples) using a white light transilluminator or a UV transilluminator with a white light conversion screen, an ultraviolet blocking cover and by wearing ultraviolet protections eyewear.

When the transfer of DNA to the membrane was finished, the membrane was introduced into a Stratalinker^®^ UV crosslinker and the Autocrosslink setting was run to fix DNA to the membrane. After this step, the blocking of the membrane can be done or the membrane can be stored in plastic cover sheets at −20°C for further use.

#### Blocking of the membrane

A TBS 10X solution (500mM Tris/Cl, 1.5 M NaCl, pH 7.5) was prepared. 30.25g of Tris/Cl, and 43.8 g of NaCl were added to a glass laboratory bottle, then 400 mL of MilliQ water was added, pH was adjusted to 7.5 and 500 mL of water were added to a final NaCl concentration of 500 mM. After this, a solution of 1×TBS-0.05% Tween20 can be prepared. The preparation consists of 100 mL of 10×TBS pH 7.5 and 0.5 mL of Tween20 added to 900 mL of Milli-Q ultrapure water.

The blocking solution was prepared with 1×TBS-0.05% Tween20 and 5% powdered milk (2.5 g of powdered milk for 50 mL of 1×TBS-0.05% Tween20).

Blocking of the membrane was done during 1h at 37°C under saturation solution and elliptical agitation.

#### Preparation of primary antibody solution and incubation

1/500 dilutions of the anti-5mC antibody (Abcam, Cat. No. ab73938, Lot: GR278832-3) were produced in blocking solution and the membrane was incubated in this solution under elliptical agitation during 1h30 at room temperature. After that, the membrane was washed three times with TBS-Tween20 for10 minutes under elliptical agitation.

#### Preparation of secondary antibody solution and incubation

1 μl of HRP-conjugated Goat anti-mouse IgG secondary antibody (Agrisera, Ref. AS11 1772, Lot: 1612) was diluted 500-fold in blocking solution.

Incubation of the membrane under elliptical agitation was done during 1h10 at room temperature. Then the antibody was removed by washing the membrane 3 times for 10 minutes in elliptical agitation

#### Reading of the signa

SuperSignal™ West Pico chemiluminescent (Thermo Fisher Scientific, Cat. No. 34580) substrate was used to detect signal intensity, a solution of 750μl of Luminol/Enhancer and 750μl of Stable Peroxide was prepared for each membrane. The solution was added to the membrane to cover it completely, then the membrane was placed on a glass plate and introduce it to the ChemiDoc™ MP Imaging System. Luminescence was captured with a CDD camera with an exposure time of 200 secs (1 photo each 10 sec) and quantified with the Image Lab™ 5.1 software. In order to determine the appropriate exposure time, the membrane was exposed to 40 secs, as DNA spots appeared too light, exposure was incremented to 3 minutes and photos were taken each 10 secs. All images were saved and the most concentrated spot in the membrane was identified, the selected exposure time was the exposure time below the point of saturation of this spot, detected by the software Image Lab (highlighting in red the saturated pixels). Normalization of the measures was performed by dividing the mean spot densitometry by the input sample DNA in ng, this provides a densitometry value per ng of DNA.

## Author Contributions and Notes

C.G and C.C. designed research, N. L. and S. D. P. performed research, N. L analyzed data; and N.L., C.G., C.C. and S.D.P. wrote the paper. The authors declare no conflict of interest.

## Acknowledgments

The authors are grateful to M. Fallet, R.Galinier, and J.F.Allienne for support during the optimization of the protocol and to N. Arancibia for breeding of mollusks. The work received funding through the Wellcome Trust strategic award 107475/Z/15/Z and through a Ph.D. grant to NL from the Region Occitanie (EPIPARA project) and the University of Perpignan Via Domitia.

## References

1. Colot, V. and Rossignol, J. L. (1999) ‘Eukaryotic DNA methylation as an evolutionary device’, Bioessays, 21(5), pp. 402–11.

2. de Lorgeril, J., Lucasson, A., Petton, B., Toulza, E., Montagnani, C., Clerissi, C., Vidal-Dupiol, J., Chaparro, C., Galinier, R., Escoubas, J.-M., Haffner, P., Dégremont, L., Charrière, G. M., Lafont, M., Delort, A., Vergnes, A., Chiarello, M., Faury, N., Rubio, T., Leroy, M. A., Pérignon, A., Régler, D., Morga, B., Alunno-Bruscia, M., Boudry, P., Le Roux, F., Destoumieux-Garzόn, D., Gueguen, Y. and Mitta, G. (2018) ‘Immune-suppression by OsHV-1 viral infection causes fatal bacteraemia in Pacific oysters’, Nature Communications, 9(1), pp. 4215.

3. Diala, E. S. and Hoffman, R. M. (1982) ‘Hypomethylation of HeLa cell DNA and the absence of 5-methylcytosine in SV40 and adenovirus (type 2) DNA: analysis by HPLC’, BiochemBiophys Res Commun, 107(1), pp. 19–26.

4. Fneich, S., Dheilly, N., Adema, C., Rognon, A., Reichelt, M., Bulla, J., Grunau, C. and Cosseau, C. (2013) ‘5-methyl-cytosine and 5-hydroxy-methyl-cytosine in the genome of Biomphalaria glabrata, a snail intermediate host of Schistosoma mansoni’, Parasites & vectors, 6(1), pp. 167.

5. Fraga, M. F., Rodríguez, R. and Cañal, M. J. (2000) ‘Rapid quantification of DNA methylation by high performance capillary electrophoresis’, Electrophoresis, 21(14), pp. 2990–4.

6. Frommer, M., McDonald, L. E., Millar, D. S., Collis, C. M., Watt, F., Grigg, G. W., Molloy, P. L. and Paul, C. L. (1992) ‘A genomic sequencing protocol that yields a positive display of 5- methylcytosine residues in individual DNA strands’, Proc Natl AcadSci U S A, 89(5), pp. 1827–31.

7. Geyer, K. K., Niazi, U. H., Duval, D., Cosseau, C., Tomlinson, C., Chalmers, I. W., Swain, M. T., Cutress, D. J., Bickham-Wright, U., Munshi, S. E., Grunau, C., Yoshino, T. P. and Hoffmann, K. F. (2017) ‘The Biomphalaria glabrata DNA methylation machinery displays spatial tissue expression, is differentially active in distinct snail populations and is modulated by interactions with Schistosoma mansoni’, PLoSNegl Trop Dis, 11(5), pp. e0005246.

8. Harrison, A. and Parle-McDermott, A. (2011) ‘DNA methylation: a timeline of methods and applications’, Front Genet, 2, pp. 74.

9. Hendrich, B. and Tweedie, S. (2003) ‘The methyl-CpG binding domain and the evolving role of DNA methylation in animals’, Trends Genet, 19(5), pp. 269–77.

10. Karimi, M., Johansson, S., Stach, D., Corcoran, M., Grandér, D., Schalling, M., Bakalkin, G., Lyko, F., Larsson, C. and Ekström, T. J. (2006) ‘LUMA (LUminometric Methylation Assay) - a high throughput method to the analysis of genomic DNA methylation’, Exp Cell Res, 312(11), pp. 1989–95.

11. Kuo, K. C., McCune, R. A., Gehrke, C. W., Midgett, R. and Ehrlich, M. (1980) ‘Quantitative reversed-phase high performance liquid chromatographic determination of major and modified deoxyribonucleosides in DNA’, Nucleic Acids Res, 8(20), pp. 4763–76.

12. Kurdyukov, S. and Bullock, M. (2016) ‘DNA Methylation Analysis: Choosing the Right Method’, Biology (Basel), 5(1).

13. Laird, P. W. (2010) ‘Principles and challenges of genome-wide DNA methylation analysis’, Nature Reviews Genetics, 11(3), pp. 191.

14. Li, W. (2011) ‘On parameters of the human genome’, J TheorBiol, 288, pp. 92–104.

15. Li, Y. and Tollefsbol, T. O. (2011) ‘DNA methylation detection: bisulfite genomic sequencing analysis’, Methods MolBiol, 791, pp. 11–21.

16. Nicoglou, A. and Merlin, F. (2017) ‘Epigenetics: A way to bridge the gap between biological fields’, Stud HistPhilosBiol Biomed Sci, 66, pp. 73–82.

17. Sant, K. E., Nahar, M. S. and Dolinoy, D. C. (2012) ‘DNA methylation screening and analysis’, Methods MolBiol, 889, pp. 385–406.

18. Sarda, S., Zeng, J., Hunt, B. G. and Soojin, V. Y. (2012) ‘The evolution of invertebrate gene body methylation’, Molecular biology and evolution, 29(8), pp. 1907–1916.

19. Schumacher, A., Kapranov, P., Kaminsky, Z., Flanagan, J., Assadzadeh, A., Yau, P., Virtanen, C., Winegarden, N., Cheng, J., Gingeras, T. and Petronis, A. (2006) ‘Microarray-based DNA methylation profiling: technology and applications’, Nucleic Acids Res, 34(2), pp. 528–42.

20. Song, L., James, S. R., Kazim, L. and Karpf, A. R. (2005) ‘Specific method for the determination of genomic DNA methylation by liquid chromatography-electrospray ionization tandem mass spectrometry’, Anal Chem, 77(2), pp. 504–10.

21. Veilleux, C., Bernardino, J., Gibaud, A., Niveleau, A., Malfoy, B., Dutrillaux, B. and Bourgeois, C. A. (1995) ‘[Changes in methylation of tumor cells: a new in situ quantitative approach on interphase nuclei and chromosomes]’, Bull Cancer, 82(11), pp. 939–45.

22. Vucic, E. A., Wilson, I. M., Campbell, J. M. and Lam, W. L. (2009) ‘Methylation analysis by DNA immunoprecipitation (MeDIP)’, Methods MolBiol, 556, pp. 141–53.

23. Walker, A. J. (2011) ‘Insights into the functional biology of schistosomes’, Parasites & vectors, 4(1), pp. 203.

24. Weber, M., Davies, J. J., Wittig, D., Oakeley, E. J., Haase, M., Lam, W. L. and Schübeler, D. (2005) ‘Chromosome-wide and promoter-specific analyses identify sites of differential DNA methylation in normal and transformed human cells’, Nat Genet, 37(8), pp. 853–62.

